# Determination of secretory granule maturation times in pancreatic islet beta-cells by serial block face scanning electron microscopy

**DOI:** 10.1101/865683

**Authors:** A. Rao, E.L. McBride, G. Zhang, H. Xu, T. Cai, A.L. Notkins, M.A. Aronova, R.D. Leapman

**Affiliations:** Laboratory of Cellular Imaging and Macromolecular Biophysics, National Institute of Biomedical Imaging and Bioengineering, National Institutes of Health, Bethesda, MD 20892; Experimental Medicine Section, National Institute of Dental and Craniofacial Research, National Institutes of Health, Bethesda, MD 20892

## Abstract

It is shown how serial block-face electron microscopy (SBEM) of insulin-secreting beta cells in wild-type mouse pancreatic islets of Langerhans can be used to determine maturation times of secretory granules. Although SBEM captures the beta cell structure at a snapshot in time, the observed ultrastructure can be considered representative of a dynamic equilibrium state of the cells since the pancreatic islets are maintained in culture in approximate homeostasis. It is found that 7.2±1.2% (±st. dev.) of the beta cell volume is composed of secretory granule dense-cores exhibiting angular shapes surrounded by wide (typically ≳100 nm) electron-lucent halos. These organelles are identified as mature granules that store insulin for regulated release through the plasma membrane, with a release time of 96±12 hours, as previously obtained from pulsed ^35^S-radiolabeling of cysteine and methionine. Analysis of beta cell 3D volumes reveals a subpopulation of secretory organelles without electron-lucent halos, identified as immature secretory granules. Another subpopulation of secretory granules is found with thin (typically ≲30 nm) electron-lucent halos, which are attributed to immature granules that are transforming from proinsulin to insulin by action of prohormone convertases. From the volume ratio of proinsulin in the immature granules to insulin in the mature granules, we estimate that the newly formed immature granules remain in morphologically-defined immature states for an average time of 135±14 minutes, and the immature transforming granules for an average time of 130±17 minutes.

## 1. Introduction

Although eukaryotic cellular function is determined at the molecular scale by biochemical pathways, function also depends on ultrastructural changes on the scale of subcellular organelles. Particularly important are processes involving changes in membranes that allow biological molecules to be sequestered in localized compartments and subsequently secreted through the plasma membrane. In animals, the secretion products of a cell can be targeted to nearby sites, e.g., release of neurotransmitters across a synapse, or to distant sites, e.g., release of endocrine hormones into the blood stream where they can affect other tissues and organs. Whereas elucidating cellular function at the molecular scale is mainly the realm of biochemistry and molecular and structural biology, it is possible to improve our understanding of cellular function on a coarser length scale with the new generation of 3D electron microscopy (EM) techniques that are applicable to entire eukaryotic cells, as well as tissues. These methods bridge the resolution gap between the nanometer scale of macromolecular assemblies accessible by cryo-electron microscopy, and the submicrometer scale accessible by fluorescent optical imaging. They include two commercially available techniques: (i) serial block-face EM (SBEM) (Denk and Horstmann, 2004; Briggman et al., 2011; Helmstaedter et al., 2013; Pfeifer et al., 2015; Shomorony et al, 2015; Deerinck et al., 2018; McBride et al., 2018; He et al, 2018), which is the one that we employ in the present study, and (ii) focused ion beam scanning electron microscopy (FIB-SEM) (Hekking et al, 2009; Bennett et al., 2009; Drobne, 2013; Narayan and Subramaniam, 2015; Glancy et al., 2018). FIB-SEM has the advantage of providing higher spatial resolution but SBEM can image much larger volumes. Since preparation of cells and tissues for electron microscopy requires fixation, either by chemical cross-linkers or by rapid freezing, cellular ultrastructure can only be captured at snapshots in time. Our aim here is to show that when a biological system is in a state of dynamic equilibrium, i.e., homeostasis, the observed ultrastructure can inform us about the behavior of these dynamic processes, if additional time-dependent biochemical data is available.

The present study focuses on mouse pancreatic islets of Langerhans — micro-organs that contain several cell types giving rise to the endocrine function of the pancreas. Mouse models have been crucial in enabling researchers to study the physiology and pathology of islet function in the laboratory. However, despite a large body of previous work, there is still much that is unknown about the structural basis of islet function (Brissova et al., 2005; Cabrera et al, 2006; Kim et al., 2009; Steiner et al., 2010; Hoang et al., 2016). Like human islets, mouse islets contain five types of hormone-secreting cells: glucagon-secreting alpha cells, insulin-secreting beta cells, pancreatic polypeptide-secreting gamma cells, somatostatin-secreting delta cells, and ghrelin-secreting epsilon cells. Beta cells are of special importance since their malfunction or destruction in humans prevents secretion of sufficient insulin causing diabetes (type 1 and type 2), which result in major morbidity and shortened lifespan throughout the world. Insulin, a small 51-residue protein with a molecular weight of 5.8 kDa, is released from islets through the hepatic vein, and functions to control elevated glucose levels in the blood. Insulin acts by increasing uptake of glucose through specific types of insulin-dependent glucose transporters in cells throughout the body, particularly muscle and fat cells, thus facilitating conversion of glucose to glycogen.

The mouse has several thousand islets each of diameter 100-200 µm, distributed throughout the pancreas and constituting 1-2% of the total pancreatic mass. Each islet contains approximately 1,000 beta cells, and each beta cell has approximately 10,000 secretory granules (Rorsman and Renström, 2003), which appear in electron micrographs as membrane-bound organelles of diameter 200 to 400 nm with dense cores surrounded by wide halos that are empty of stain. Some of the granule cores appear angular in shape consistent with the formation of insulin crystals. It is known from other experiments that the insulin can be packaged in crystalline form after association with zinc (Foster et al., 1993; Goping et al., 2003; Li, 2014). In mice, 65-80% of the islet cells are beta cells, and 10-25% alpha cells, with only 5-10% of the cells belonging to the other three cell types. Islets also contain intricate networks of fenestrated capillaries that permeate through their interiors with pericapillary space organized so that each beta cell makes direct contact with the blood plasma (Bonner-Weir, 1993). After isolation and transfer to culture medium, islets have been shown to maintain their physiological state including response to elevated glucose, although the capillaries are no longer intact.

Previous work has suggested that a small fraction of the beta cell’s insulin is released without being stored in secretory granules via the constitutive pathway (Lara-Lemus et al., 2006; Boland et al., 2017), but by far the largest component of insulin is packaged into secretory granules and released through the regulatory pathway, typically ∼99% (Halban, 1991). It has been proposed that newly formed secretory granules are transported along microtubules from the trans-Golgi network towards the cell periphery, where they are captured in the F-actin-rich cortex where they are immobilized and mature before eventual release (Rudolf et al., 2001).

### 1.1. Use of serial block-face EM to visualize 3D ultrastructure of entire beta cells

Here, we use the technique of serial block face electron microscopy (SBEM) to image beta cells of pancreatic islets in wild-type mice. SBEM provides the 3D nanoscale architecture of entire eukaryotic cells, enabling structural biologists to visualize the totality of organelles and their connectivity, as well as to obtain quantitative information about the numbers and sizes of subcellular compartments. In SBEM, an ultramicrotome built into the specimen chamber of a scanning electron microscope (SEM) allows imaging of large volumes of tissues stained with heavy atoms and embedded in plastic, with a voxel size of ∼5 nm in the plane of the block face, and ∼25 to ∼50 nm in the perpendicular direction, as limited by the minimum slice thickness that the ultramicrotome can remove (Fig. 1A). In an automated process, the face of the sample block is imaged by detecting backscattered electrons, which provides strong contrast from the stain. The block is raised by some set amount (e.g., 25–50 nm), and is then cut by the ultramicrotome to expose a new surface. Repetition of this process enables sequential acquisition of several hundred block-face images, each of which corresponds to an orthoslice through the 3D volume (Fig. 1B).

**Figure 1.**
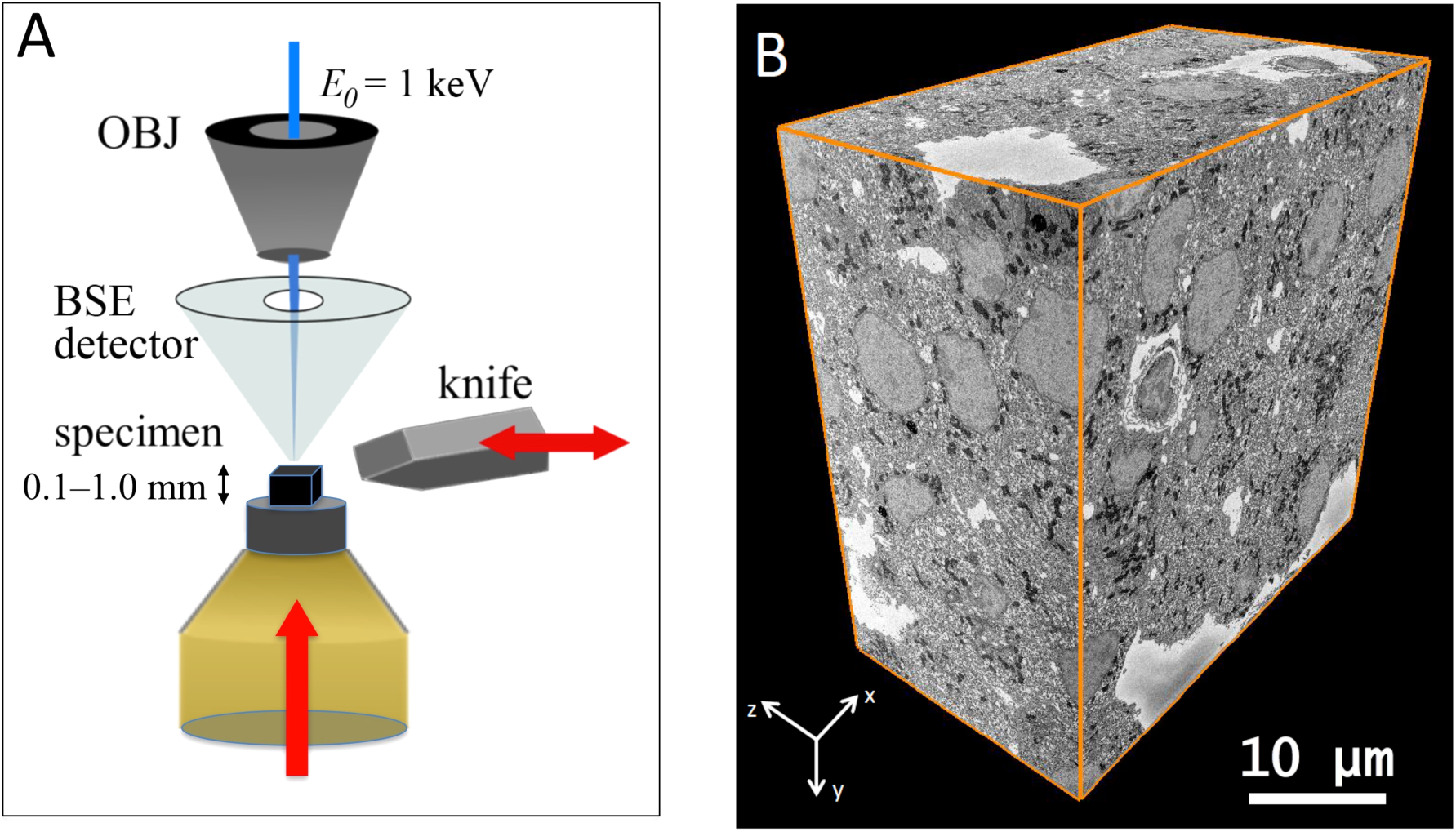
(A) Schematic diagram of Serial Block-Face Scanning Electron Microscopy (SBF-SEM). After imaging with a back-scattered electron detector (BSD), a diamond knife cuts a thin slice from the face of the block. The newly exposed block face is then imaged, and the knife cuts again. This repetition of cutting and imaging allows one to build up a set of images, which represent the volume of the object imaged with the electron beam. (B) A part of wild type mouse islet 3D data set obtained by SBF-SEM. This data measures 42.6 μm × 43.5 μm × 23.9 μm in *x y* and *z*, respectively.

In an earlier study, we have used SBEM to quantify distributions of dense-core granules in beta cells of mouse islets. Here, we use this technique to obtain quantitative measurements on immature secretory granules in beta cells. Our results thus provide an estimate of the time taken for newly formed granules containing proinsulin to mature into dense-core insulin granules that are stored intracellularly for eventual regulated release of the hormone through the plasma membrane. Such information cannot be obtained from conventional transmission electron microscopy (TEM) of thin plastic sections because organelles cannot be observed in their entirety. For example, it is impossible to know from the observation of an empty membrane-bound structure in a TEM image whether that structure corresponds to a section through an empty vesicle, or whether it corresponds to a section through a dense-core granule in which the dense core happens to be absent. Such ambiguities are immediately resolved if the full 3D structure is available, as is the case for images acquired in SBEM. Similarly, it is difficult to establish from TEM images whether the diameters of secretory granules lie within a narrow range or whether their diameters form a wider distribution; again, such uncertainties are avoided by analysis of SBEM data. Furthermore, in TEM images, the immature granules are found to be quite sparsely distributed and it can be difficult to assess their overall frequency in the cell or even to know whether all beta cells contain those structures. SBEM offers a considerable advantage in this regard because entire cells can be visualized and rare structures quantified. As we have shown previously, it is also possible to perform faster analyses of SBEM data through a combination of 2-D and 3-D measurements, by making use of stereological techniques, which we employ here too.

### 1.2. Ultrastructure of pancreatic beta cells and a model of insulin secretion

Careful examination of cellular ultrastructure obtained from transmission electron microscopy (TEM) or SBEM of fixed, stained, and embedded mouse pancreatic islets of Langerhans reveals that not all secretory granules in beta cells have densely stained cores surrounded by clear and wide halos; a small subpopulation of granules exhibit cores that are less strongly stained and that contain much thinner halos, or no halos at all (Fig. 2). We identify these structures as immature, newly formed granules that bud off and detach from the trans-Golgi network (Furuta et al., 1998). It is well known that the immature secretory granules contain proinsulin, which is a single peptide of molecular mass 8.93±0.01 kDa composed of a C-chain containing 29 residues (mouse insulin 1) or 31 residues (mouse insulin 2) connecting an A-chain containing 21 residues to a B-chain containing 30 residues (Imai, 2015). Proinsulin is produced from another precursor protein, pre-proinsulin, which is synthesized in the endoplasmic reticulum and contains additional amino-acids in the form of a signal peptide that is cleaved off before reaching the Golgi apparatus. Examples of immature granules in the region of the trans-Golgi network are shown in Fig. 3A. After being sorted into immature secretory granules, the C-peptide is cleaved from proinsulin by a series of enzymes comprising the prohormone convertase (PC) family of serine proteases including PC1/3 and PC2 (Muller and Lindberg, 2000), and furin (Louagie et al., 2008), after acidification of the secretory granules from a pH of approximately 6.3 down to 5.5 at which the proteases have maximal activity (Furuta et al., 1998). Evidence for the first stages of maturation of the newly formed, proinsulin-containing, immature granules is the occurrence of a subset of granules displaying narrow halos, which appear to consist of a distinct subpopulation in which prohormone convertases have begun to produce insulin by cleavage of C-chain of proinsulin (Fig. 3B). After cleavage of the C-chain, the A-chain and B-chain of insulin remain tethered to each other through disulfide bonds in the form of the functional hormone insulin, which is then able to crystallize as hexamers in the presence of divalent zinc ions, resulting in granules containing dense cores with angular morphology. The relative insolubility of the dense crystalline cores of mature secretory granules makes this form of insulin suitable for longer term storage in the beta cells. Segmentation of the cores and outer membranes of mature and immature transforming secretory granules are shown in Figs. 4A and 4B, respectively; the corresponding 3-D surface-rendered views of these mature and immature transforming granule are represented in Figs. 4C and 4D, respectively. It is evident that the mature granule has an asymmetrical halo that extends approximately 200 nm from the core at its widest point, whereas the halo of the immature transforming granule has an approximately constant width of about 30 nm.

**Figure 2.**
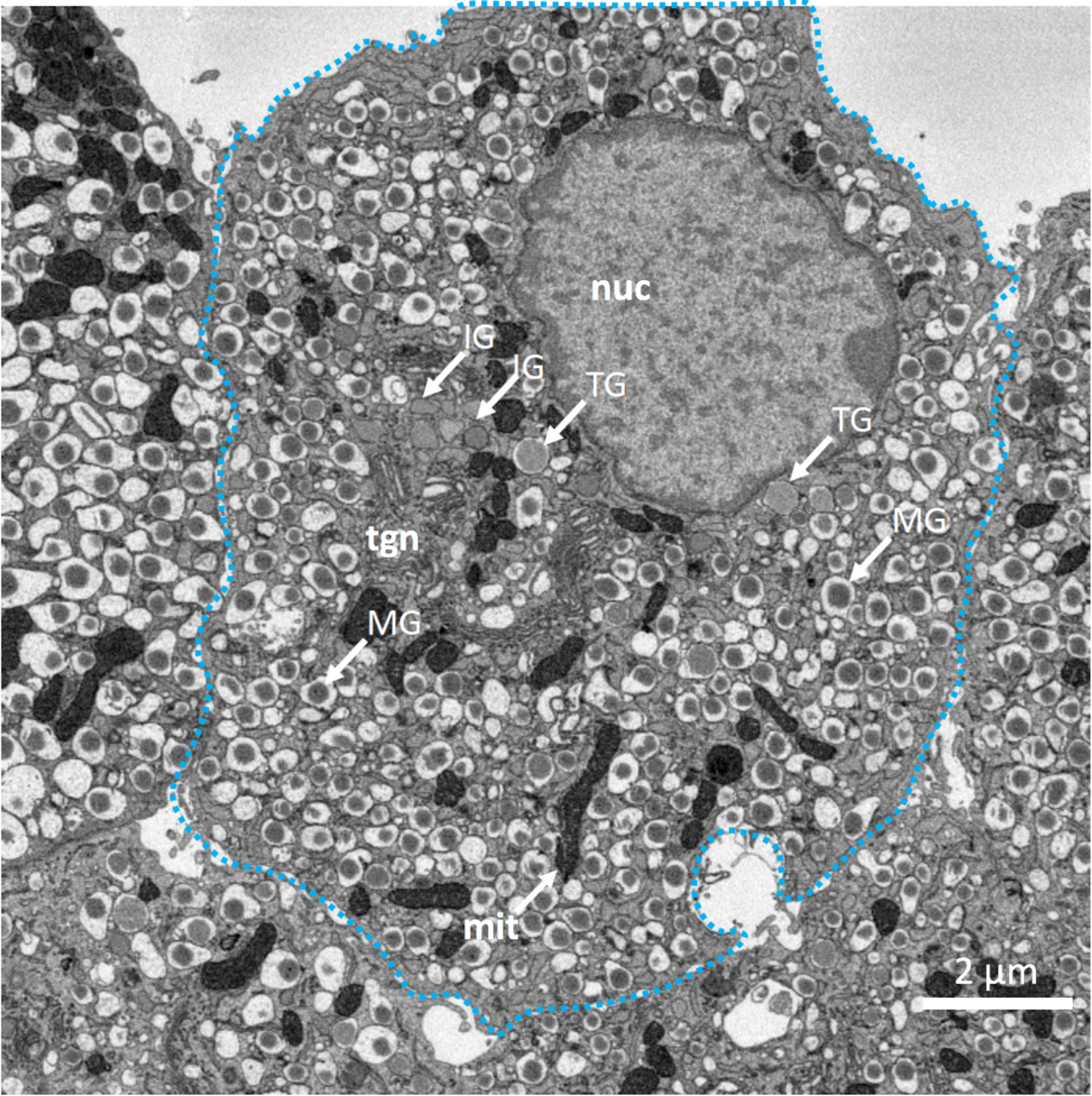
SBEM backscattered electron image of section through pancreatic beta cell indicating morphologies of mature secretory granules (MG), immature secretory granules (IG), and transforming secretory granules (TG); mature granules contain cores of condensed insulin with angular sides and large surrounding halos. Immature granules are situated mainly near the trans-Golgi network (tgn) and have no halos. Several of the immature granules are near nucleus (nuc), while others are interspersed with mitochondria (mit). The transforming granules form a distinct population with very narrow halos.

**Figure 3.**
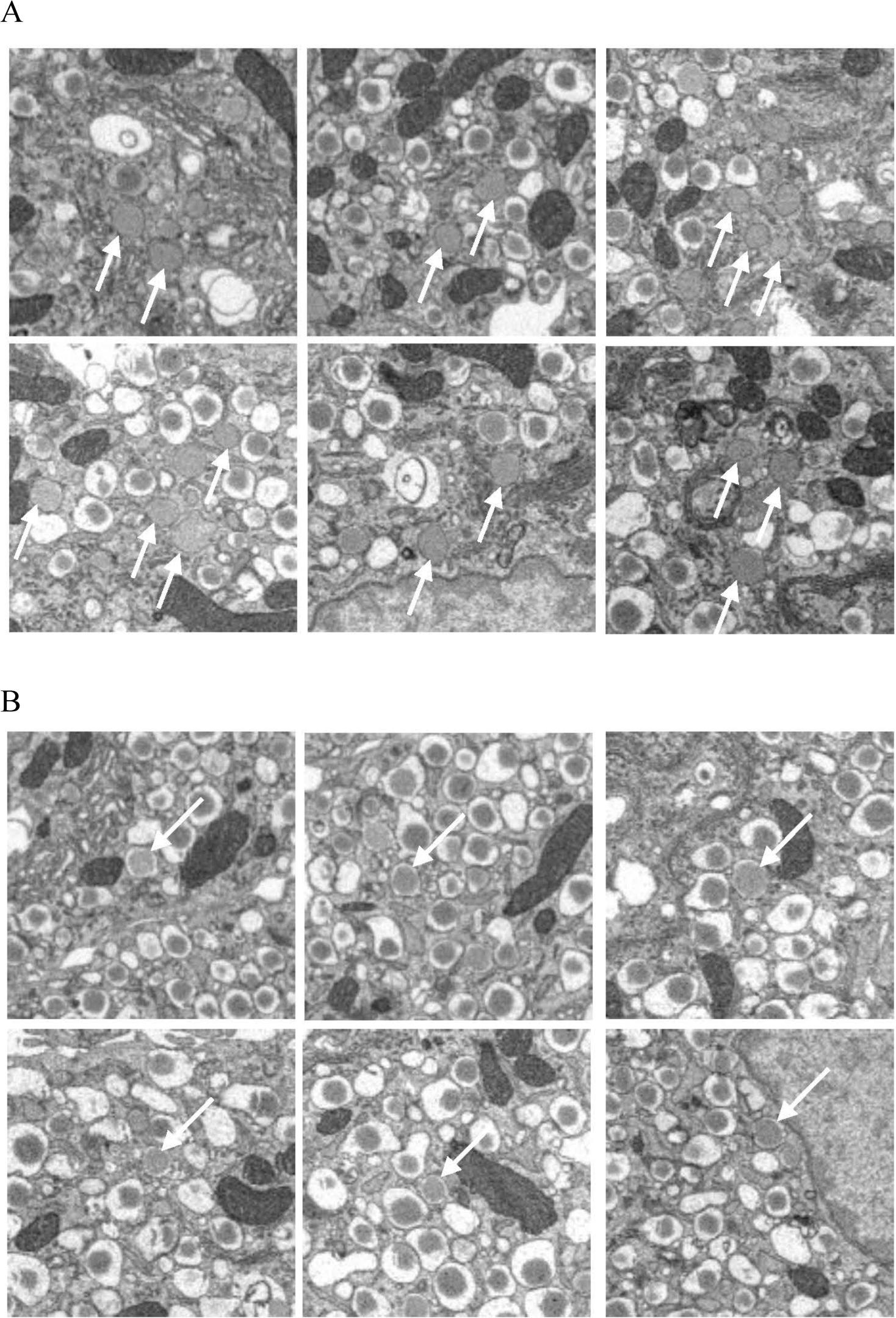
(A) Six representative areas of beta cells in the vicinity of the trans-Golgi network, showing immature granules that have no halos. (B) Six representative areas of beta cells, showing transforming immature granules with narrow halos. Full width of each image is 4 µm.

**Figure 4.**
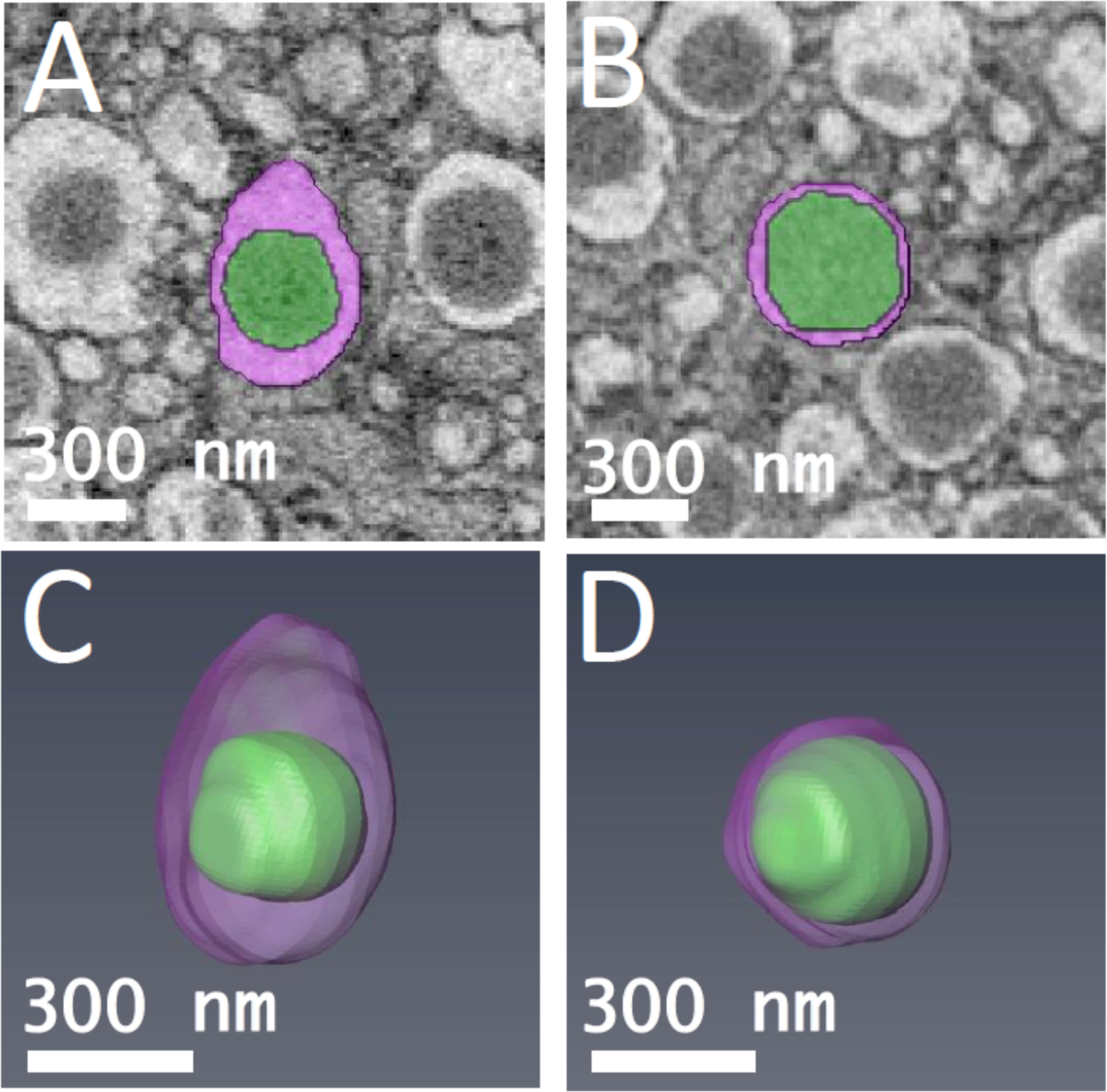
Segmented orthoslices of (A) mature β-cell secretory granule, and (B) transforming immature β-cell secretory granule. The granule cores are labeled in green, and the surrounding halos are labeled in purple. 3-D models of (C) mature β-cell secretory granule shown in (A), and (D) transforming immature β-cell secretory granule shown in (B). The ratio of core volume to total vesicle volume was measured for both granules.

In addition to storing insulin, beta cell secretory granules also store and co-secrete the satiation hormone amylin or amyloid islet polypeptide, which contains 37 amino acids; the prohormone proamylin contains 67 amino acids and is converted like proinsulin to the hormone in the secretory granule. Amylin is present at only around 1% of concentration of insulin and is believed to be localized in the secretory granule halo along with the soluble cleaved C-chain of insulin (Lutz and Meyer, 2015; Karlsson et al., 1996). Since the relative concentration of amylin is low, we neglect its effect in the morphological analysis of insulin and proinsulin in the beta cell granules.

A simplified description of the maturation of beta cell secretory granules is summarized in Fig. 5, which includes our observations of granules without halos situated close to the trans-Golgi network, and granules with narrow halos where we believe proinsulin is starting to undergo conversion to insulin after acidification and subsequent action of prohormone convertases. The diagram in Fig. 5 also indicates the production of mature granules with crystalline dense cores and wide halos, which eventually fuse with the plasma membrane to secrete insulin.

**Figure 5.**
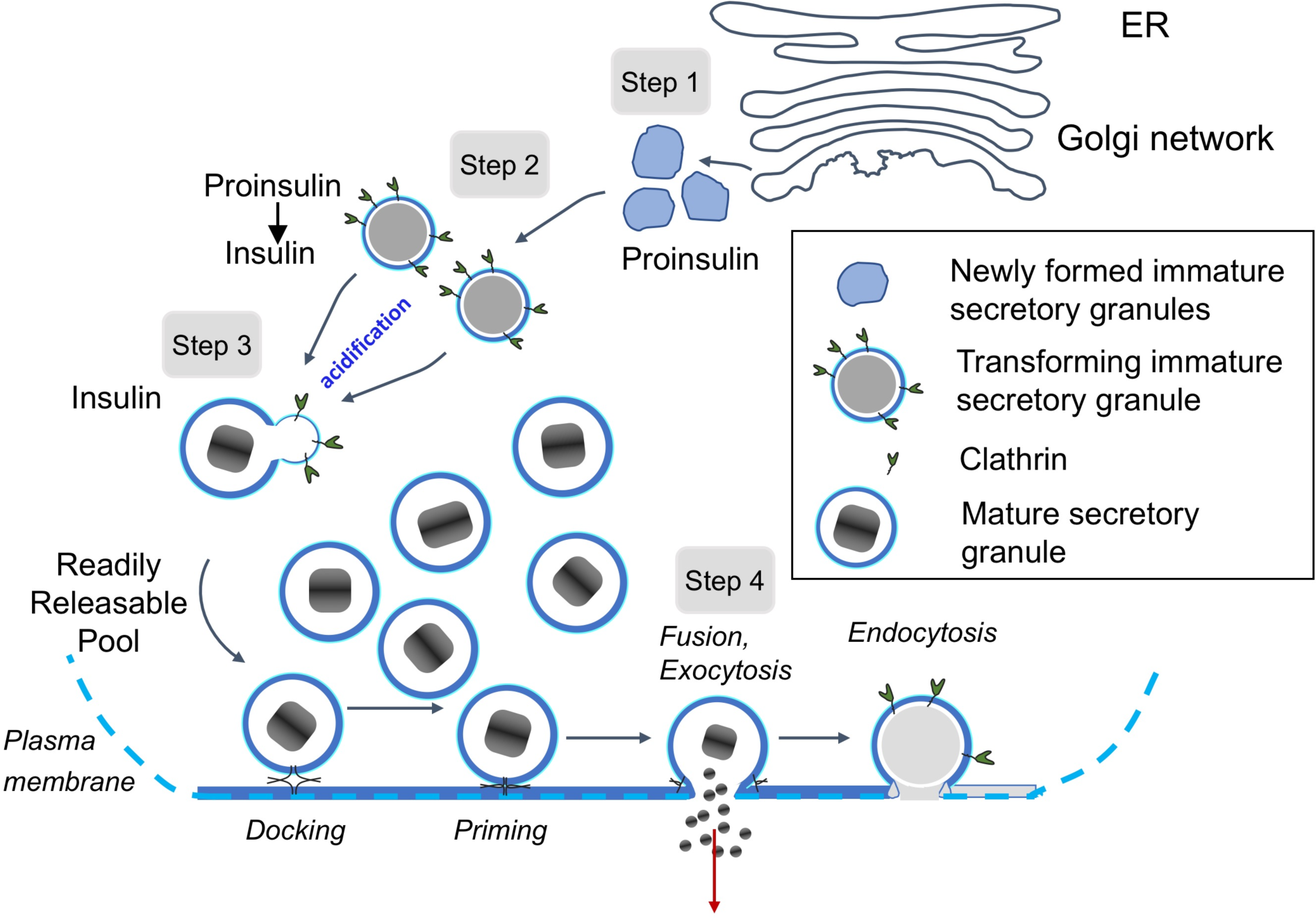
Simplified diagram depicts maturation and secretory pathway of insulin secretory granules in wild-type mouse pancreatic islet beta cells. In Step 1, proinsulin is packaged into vesicles that bud off from trans-Golgi network as immature secretory granules. In Step 2, the pH of immature granules is lowered facilitating action of prohormone convertases, which begin to cleave the soluble C-peptide chain from proinsulin, as evidenced by the presence of clear, narrow halos. In Step 3, action of the prohormone convertases proceeds rapidly (as evident by relatively few granules with narrow halos), and the secretary granules become mature, displaying crystalline cores of insulin. In Step 4, mature granules fuse with the plasma membrane, release insulin to the extracellular space, and the membrane is recycled through clathrin-mediated endocytosis. The granules are sorted for biphasic release, in which granules fuse to the membrane and release their contents.

## 2. Materials and Methods

### 2.1 Sample preparation

Pancreatic islets from 2–3-months-old wild type male mice were isolated as described previously (West et al., 2010; Holcomb et al., 2013). Mice were anesthetized by intraperitoneal injection with ketamine (50 mg/kg), and collagenase solution (Sigma, USA) was injected into the bile duct to inflate the pancreas. After digestion, islets were manually selected and washed in Krebs–Ringer HEPES buffer, and cultured overnight in RPMI-1640 medium (Invitrogen, USA). All procedures were approved by the NIDCR Institutional Animal Care and Use Committee. The islet fixation procedure proceeded as previously described (Pfeifer et al., 2015). After aldehyde fixation, the islets were fixed in reduced osmium. Samples were then processed with Walton’s lead aspartate staining (Walton, 1979) by submerging the islets in a lead aspartate solution and placing them in an oven for 30 min. Then, the samples were dehydrated and embedded in Epon-Araldite by following standard protocols. A block of resin-embedded islets was first mounted on an empty resin block to be trimmed under the microtome to select approximately one islet. Once the top of the stained islet was exposed, the islet was re-mounted, exposed-side down, on an aluminum specimen pin (Gatan Inc., USA) using CircuitWorks conductive epoxy (CW2400). Each re-mounted sample was then trimmed again to expose the other side of the islet. The block was then sputter-coated with a 40 nm-thick layer of gold to increase electrical conductivity and to prevent specimen charging.

### 2.2. Serial block face scanning electron microscopy

The resin-embedded, gold-coated blocks were imaged using a Gatan 3View serial block-face imaging system mounted in the specimen chamber of a Zeiss SIGMA-VP SEM. A total of five blocks (one islet in each), taken from three wild-type mice, were imaged to collect the data used in this study. The microscope was operated at high vacuum, with accelerating voltage ranging from 1.3-1.5 kV, using a 30 μm condenser aperture. Images were collected at an intermediate magnification, with a pixel size of 11.1 nm and a dwell time of 2.0 μs/pixel. The size of each image collected was 4,000 × 4,000 pixels in the x-y plane. The thickness of each slice in z-direction was 50 nm. Data sets were collected as stacks of 500 images, resulting in data volumes of 4.93 × 10^4^ μm^3^. The 3D image stacks were aligned using Digital Micrograph software (Gatan Inc., USA). Visualization and analysis of the data were performed using Digital Micrograph and Amira (FEI Inc., U.S.A.), as well as the NIH ImageJ software (Schneider et al., 2012).

### 2.3. Determination of numbers of moles of proinsulin and insulin per beta cell from image stacks

To estimate the lifetime of the newly formed proinsulin granules, and time taken for the proinsulin granules to transform into mature insulin granules, it is insufficient to measure the numbers of secretory granules in these different states, since it is possible that granules fuse with each other or split apart during the maturation process. Instead, it is preferable to estimate the number of moles of proinsulin per beta cell in these two cellular compartments, as well as the number of moles of mature insulin per cell.

The volume fraction of newly formed immature secretory granule cores *f*_*im*_ can be written as:

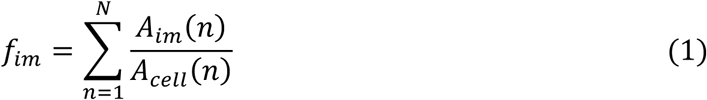

where *A*_*im*_(*n*) is the summed area of all the immature secretory granule cores in slice *n* of the image stack, and *A*_*cell*_(*n*) is the area of the cell in slice *n*, and the volume fraction of immature granules that are transforming into mature granules, defined by narrow halos, *f*_*tr*_ can be written as:

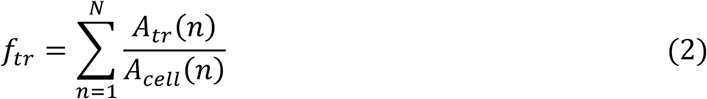

where *A*_*tr*_(*n*) is the summed area of all the transforming secretory granule cores in slice *n*. The volume fraction of mature secretory granules *f*_*mat*_ can be written as:

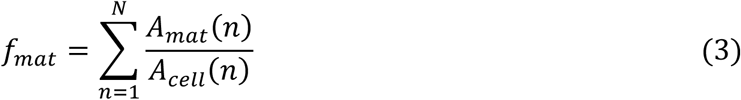

where *A*_*mat*_(*n*) is the summed area of all the mature secretory granule cores in slice *n* of the image. If the mean volume of the beta cell is *V*_*cell*_, the density of proinsulin is *ρ*_*proins*_, the density of insulin is *ρ*_*ins*_, the molecular weight of proinsulin is *M*_*proins*_, and the molecular weight of insulin is *M*_*ins*_, then the number of moles *Q*_*im*−*proins*_ of proinsulin per beta cell contained in immature granules is given by:

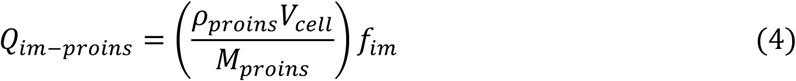

and the number of moles *Q*_*tr*−*proins*_ of proinsulin per beta cell contained in transforming secretory granules is given by:

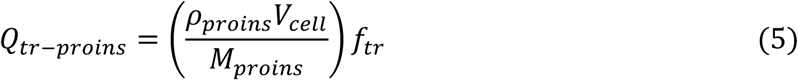

and the number of moles *Q*_*ins*_ of insulin per beta cell contained in mature secretory granules is given by:

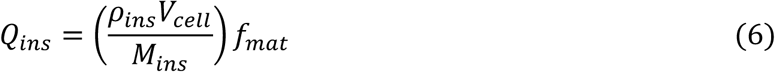

### 2.4. Measurement of half-life for insulin secretion by radioimmunoassay (RIA)

An important parameter that we use in estimating the maturation time of secretory granules is the half-life for insulin secretion from the beta cells. We have previously measured the insulin half-life using radioimmunoassay by pulse-chasing isolated wild-type mouse islets with ^35^S-labeled amino acids (Cai et al. 2011). Isolated wild type islets were pulsed with [^35^S]methionine and [^35^S]cysteine. In brief, groups of 25 islets on a 62 μm monofilament nylon mesh inside 13 mm Swinnex chambers (Millipore, Bedford, MA, USA) were perfused at 0.5 ml/min with KRB buffer containing 2.8 or 16.7 mmol/l glucose. Aliquots of perfusate (0.5 ml) were collected at different times over 1 h and stored at −80°C for measurements (Insulin RIA kit, Linco, St Charles, MO, USA). Insulin content in acidic alcohol-extracted islets was determined by RIA. The half-life of insulin was determined to be *t*_*R*_ = 96 ± 12 h, which is in the range reported in the literature for other WT mice (72–120 h) [14].

### 2.5. Method for estimating life-times of immature secretory granules in beta cells

Although our analysis has thus far only provided a view of granule morphology in snapshots of time, with certain assumptions, we can derive some dynamic information about the lifetime of the immature secretory granules located in the region of the trans-Golgi network, as well as the lifetime for the transforming secretory granules characterized by thin halos. The first assumption is that beta cells are in a steady state of dynamic equilibrium, in which granules evolve from one morphological phase to another, beginning with the immature granules that bud off the trans-Golgi network, their movement throughout the cell as transforming secretory granules with thin halos, and their maturation into dense core granules with wide halos, and finally their fusion with the plasma membrane with release of insulin into the extracellular space. The number of moles of proinsulin or insulin contained within these different morphologically distinct granule phases must then be proportional to the duration of that phase. The second assumption is that all granules share the same pathway from the new formation of immature granules in the trans-Golgi network, to the first morphologically indicated maturation stage by action of prohormone convertases and concomitant appearance of a narrow halo following cleavage of the C-peptide, followed by appearance of fully mature crystalline-shaped dense granule cores containing insulin, and finally the secretion of insulin from the beta cell by fusion of the mature granules with the plasma membrane.

Although the release of insulin from beta cells is known to be pulsatile and biphasic (Hou et al., 2009; Rorsman and Renström, 2009; Hoange, et al., 2016), only small fractions of the beta cell’s ∼10,000 mature secretory granules release their insulin in response to high glucose (0.14% per minute and 0.05% per minute for first-phase and second-phase, respectively). This means that there remains a very large buffer capacity of stored insulin within the beta cell so that, to first order, the biphasic and pulsatile behavior of insulin release is not expected to have an immediate effect on the production of proinsulin granules or the rate at which they mature.

We consider that *Q*_*ins*_ is the number of moles of insulin that are contained in mature granules with wide halos and densely stained cores, which have an average lifetime of *τ*_*ins*_ in the cytoplasm of the beta cell; *Q*_*im*−*proins*_ is the number of moles of proinsulin that are contained in immature granules without halos, which remain in that state for time *τ*_*im*_; and *Q*_*tr*−*proins*_ is the number of moles of proinsulin that are contained in transforming granules with thin halos and lightly stained cores, which remain in that state for time *τ*_*tr*_. Then, we can write:

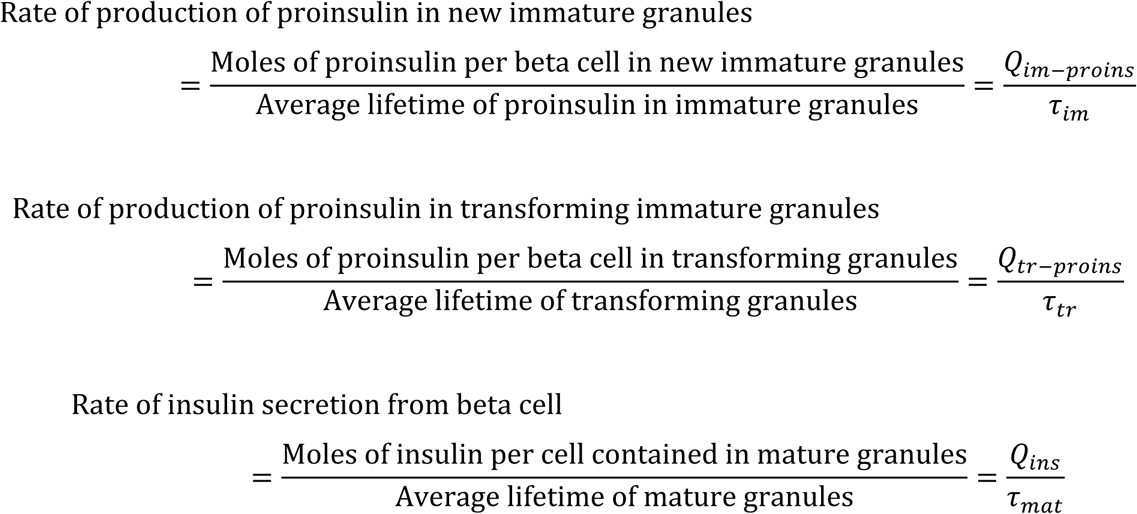

In equilibrium, the three rates must be equal so that:

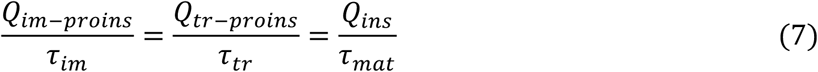

If we consider that the experimentally determined time required for the beta cell to release half of its stored insulin through fusion of mature granules with the plasma membrane is *τ*_*R*_, then the average half-life of a mature granule, *τ*_*mat*_ = *τ*_*R*_. As indicated above, an estimate of *t*_*R*_ can be obtained by labeling the insulin isotopically in the beta cells and measuring the time for the detected isotope to decrease to half of its original level.

Therefore, the average lifetime of an immature secretory granule is given by:

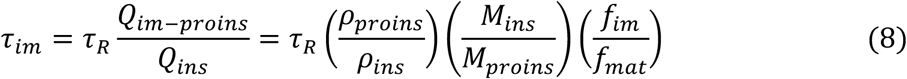

and the average lifetime of the transforming granule is given by:

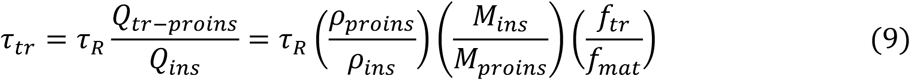

## 3. Results and Discussion

### 3.1. Measurements of volume fractions of mature insulin granule cores, immature granules cores, and transforming granule cores

Secretory granule areas were measured in block-face surfaces cut through the volumes of six beta cells from five pancreatic islets of Langerhans originating from three different mice. A complete data set from one beta cell is presented in Supplemental Movie 1. In all, we measured the areas of 8,120 mature granules in 50 images through the beta cell image stacks with block-faces selected at a spacing 1 µm, and with each image containing between about 100 and 300 mature granules, as illustrated for one beta cell in Fig. 6A, where the core areas were segmented by contouring. Since the numbers of immature granules and transforming granules were much lower than the number of mature granules, we performed measurements on 117 images from block-face surfaces through six cells, mostly sampled with a spacing of 0.5 µm, rather than 1 µm. The granules core areas from a total of 396 immature granules and a total of 397 transforming granules were segmented by contouring, as illustrated for the same beta cell in Figs. 6B and 6C, respectively. The determinations of mean fractional cell cross sectional areas, equal to the fractional cell volume, occupied by mature, immature and transforming secretory granule cores are summarized in Table 1, and presented as histograms in Figs. 7A, 7B, and 7C, respectively.

**Table 1.** Fraction of beta cell volume for different stages of secretory granules obtained from slices through cellular volumes spaced 1 µm or 0.5 µm apart in SBEM image stacks.

|  | Cores of mature secretory granules | Cores of immature secretory granules | Cores of transforming secretory granules |
| --- | --- | --- | --- |
| Fractional volume occupied by secretory granules (mean of 6 cells) | $f_{mat} = 0.072$ | $f_{im} = 0.0026$ | $f_{tr} = 0.0025$ |
| st. dev. of fractional volume occupied by secretory granule type | 0.012 | 0.0005 | 0.0007 |
| s.e.m. of fractional volume occupied by secretory granule type | 0.005 | 0.0002 | 0.0003 |
| Total number of granule type analyzed | 8,120 | 396 | 397 |
| Number block face images analyzed | 50 | 117 | 117 |

**Figure 6.**
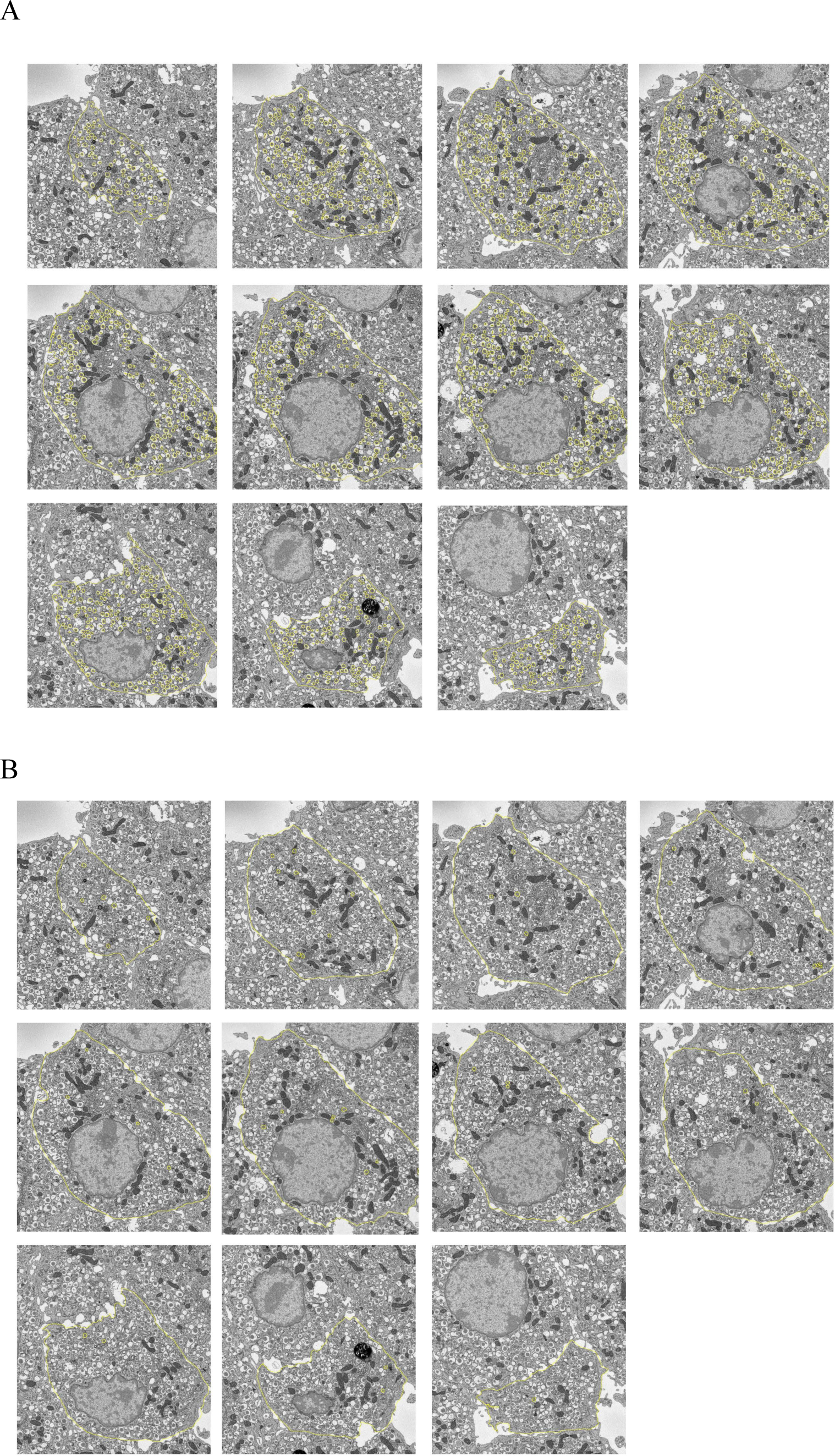
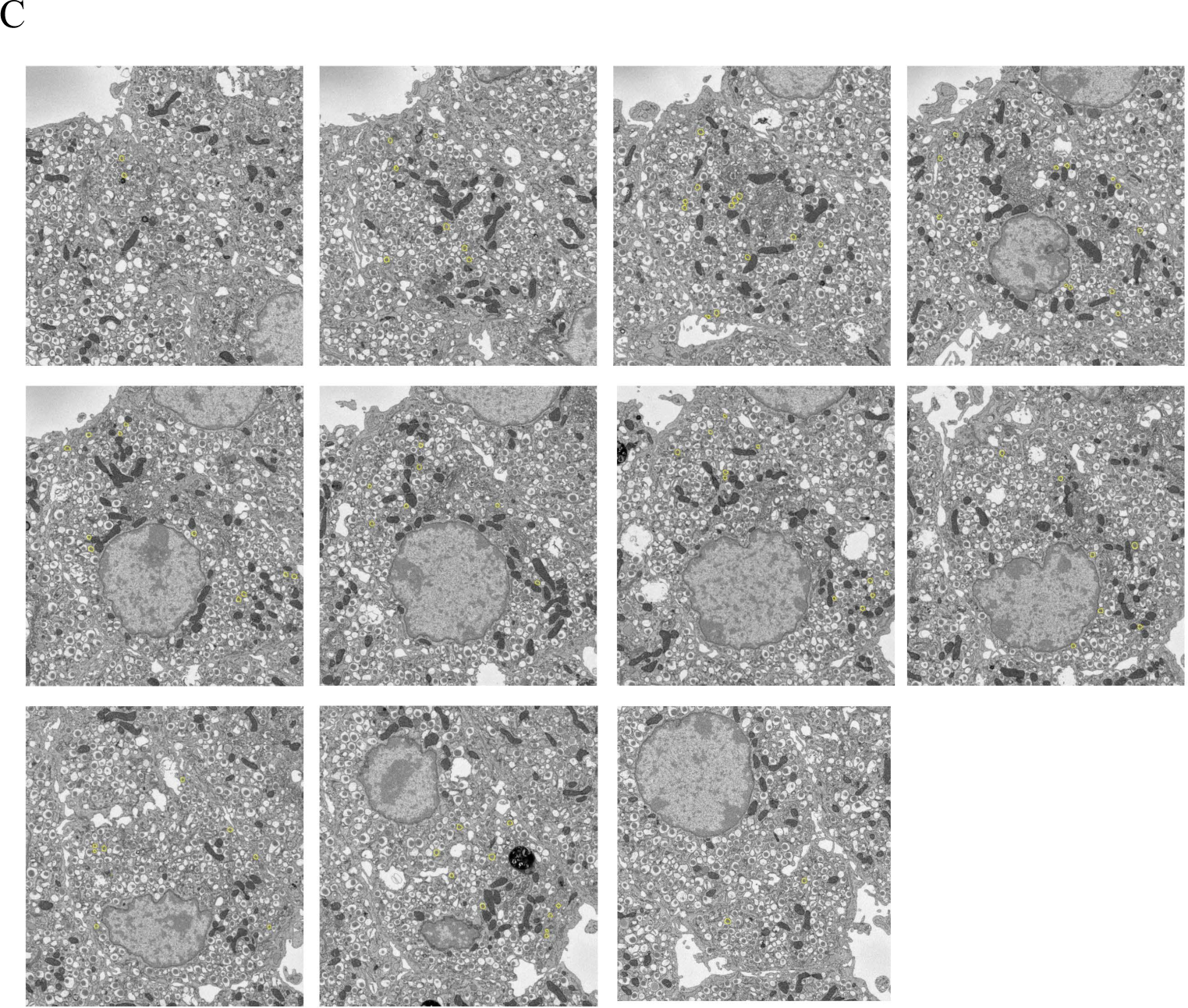
Segmentation of secretory granules in 11 slices spaced by 1 µm through a beta cell: (A) mature granules, defined by wide halos (typically ≳100 nm) and angular-shaped (crystalline) cores; (B) immature granules, most commonly located in vicinity of trans-Golgi network, and defined by absence of halos and amorphous cores; (C) transforming immature granules, defined by narrow halos (typically ≲30 nm). Cell membrane is outlined in (A) and (B). Full width of each image = 15 µm.

**Figure 7.**
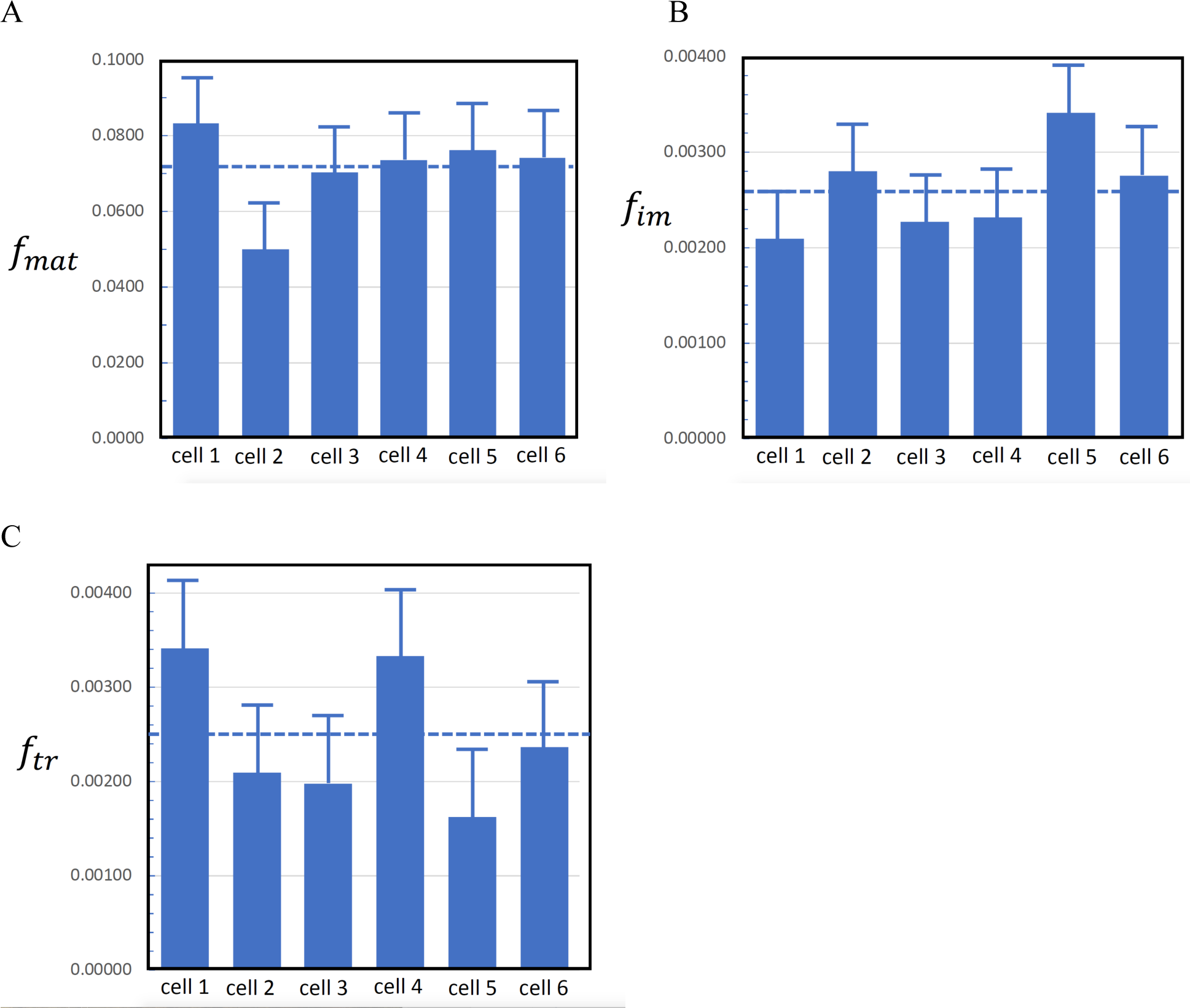
Volume fractions of secretory granule cores in six pancreatic islet β-cells (A) mature granules; (B) immature granules; (C) transforming granules. Bars represent standard deviation in the measurements.

The volume fraction of mature secretory granules in the beta cells is consistent with previous analysis to within the experimental error. In that work it was found that the area fraction of mature secretory granules cores in random slices through granule-rich regions of beta cells was 0.132±0.027 (Shomorony et al., 2015); and taking measured fraction of 0.287 of the cell volume occupied by nucleus and mitochondria, i.e., excluding secretory granules, the cellular volume fraction occupied by secretory granule cores would be (0.132 ± 0.027) × (1 − 0.287) = 0.094 ± 0.019. However, this earlier work did not consider the volume fraction of immature or transforming vesicles, nor other cellular regions that exclude secretory granules such as endoplasmic reticulum and Golgi. The measurements in Table 1 do not make assumptions about the volume of the granule-rich regions of the cell, or the regions excluded by the nucleus, mitochondria, and other organelles. Instead, the present data are based on random sections through entire cells, and measurements were made on a total of 8,120 mature secretory granule cores.

### 3.2. Ratio of proinsulin to insulin obtained from morphological measurements and comparison with biochemical measurements

From Eqs. (6)-(8), we can express the molar ratio of proinsulin to insulin in beta cells as:

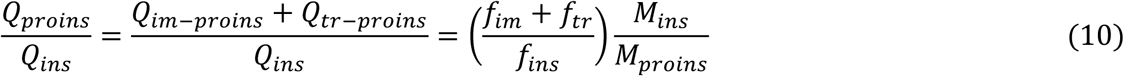

Here, we have included proinsulin in the newly formed immature secretory granules, as well as insulin in the secretory granules containing narrow halos, which are beginning to transform to the mature state. In fact, we initially believed that all beta cell immature secretory granules contained narrow halos, but further examination of the SBEM data indicated that the organelles, which bud off the trans-Golgi network, contain no halos and represent the newly formed immature granules. In Eq. 10, we have also assumed that the density of proinsulin packaged into secretory granules is approximately equal to the density of insulin, based on x-ray crystallographic data (Fullerton et al., 1970; Badger and Caspar, 1991; Badger et al., 1991).

From the known peptide sequences, it is known that the molecular mass of both mouse insulin 1 and insulin 2 is 5.80±0.01 kDa, and the molecular mass of mouse proinsulin is 8.93±0.01 kDa, prior to cleavage of the C-peptide. Then, from our measured values of *f*_*mat*_, *f*_*im*_, and *f*_*tr*_ in Table 1, we find that *Q*_*proins*_/*Q*_*ins*_ in mouse beta cells is 0.046±0.005 (± s.e.m.). This result is consistent with previous published data from rat islets, which gave a ratio of proinsulin to insulin plus proinsulin of 0.055, based on an immunoradiometric assay (Yue et al., 1979).

### 3.3. Estimated lifetimes for newly formed and transforming immature secretory granules

The fractions of summed areas of newly formed immature granule cores *f*_*im*_, transforming granule cores *f*_*tr*_, and mature granule cores *f*_*mat*_, relative to the areas enclosed by the beta cell plasma membrane were determined from SBEM image stacks by means of Eqs. 1 to 3. Mean values of the area fractions were: *f*_*im*_ = 0.0026 ± 0.0005, *f*_*tr*_ = 0.0025 ± 0.0007, *f*_*mat*_ = 0.072 ± 0.005, where the uncertainties are now expressed as standard errors rather than as standard deviation, since we believe that variations in the measured area fractions are mainly due to Gaussian statistics of structures sampled in random sections through the image stacks. These mean area fractions of the secretory granule cores are equal to the corresponding volume fractions, based on straightforward stereological considerations.

Our measurement of mean volume fraction of mature granule cores *f*_*mat*_, together with the estimated density of the crystalline insulin core *ρ*_*ins*_ = 0.47 g/cm^3^ from x-ray crystallographic data (Badger and Caspar, 1991), gives an estimate a mass of insulin per unit volume of *ρ*_*ins*_*f*_*mat*_ = 0.072 × 0.47 = 0.034 g/cm^3^. This result is reasonably consistent with previously reported data of Lacy and Richardson (1962) who found a dry mass fraction of 0.145 ± 0.015, which corresponds to 0.043 ± 0.005 grams of insulin per hydrated gram of beta cells assuming a 70% water content, and Declercq et al. (2010), who found 0.050 grams of insulin per gram of beta cells.

We can now use Eqs. 8 and 9 to estimate the lifetimes of the newly formed immature vesicles characterized by lack of halos, and the morphologically distinct transforming immature vesicles, characterized by narrow (< 30-nm wide) halos, which we believe are formed after acidification of the immature granule and action of the proenzyme convertases. It is found that the lifetime of the newly formed immature granules is *τ*_*im*_ = 135 ± 14 minutes, and the lifetime of the transforming immature granules is *τ*_*tr*_ = 130 ± 17 minutes.

### 3.4. Comparison of estimates of maturation times based on radioactivity measurements in pulse chase experiments

It is interesting to compare our estimates of maturation times for beta cell secretory granules with experiments by Rhodes et al. (1987), in which isolated rat islets were pulse-labeled with [^3^H]leucine and then incubated for a 25-minute chase period before being homogenized and fractionated to provide a purified beta granule preparation, which was re-suspended in a buffer to simulate the beta cell cytoplasm. In the presence of ATP, which activated the beta granule proton pump and lowered the granule pH, Rhodes et al. (1987) found that approximately 40% of the endogenously labeled [^3^H]proinsulin in isolated rat beta cell granules was converted to insulin in a period of τ_im_ = 120 minutes by measuring radioactivity in components isolated from high-performance liquid chromatography. These authors also monitored conversion of [^3^H]proinsulin to insulin in intact rat islets that had been pulse-labeled with [^3^H]leucine, and found that 77% of the proinsulin was converted to insulin in a period of 120 minutes.

In another study, Furuta et al. (1998) performed pulse-chase experiments on wild-type mouse islets and on islets in which PC2 convertase gene was mutated. Those authors found that in wild-type islets, approximately 90% of the proinsulin was converted to insulin in 2 hours, whereas in the mutant, the conversion time for proinsulin increased about three-fold. Furuta et al. (1998) also performed immuno-electron microscopy, and showed a large increase in the number of immature (maturing) secretory granules in the mutant islets, demonstrating the action of PC2 convertase both morphologically as well as biochemically. Our morphological determinations of granule maturations times based on the total masses of proinsulin and insulin contained in the secretory granules (135±14 minutes for the immature granule lifetime, and 130±17 minutes for the transforming granule lifetime) are therefore in agreement with these earlier pulse-chase measurements.

Our results show the consistency between three sets of independent measurements: (i) ratio of numbers of moles of proinsulin (in the cores of immature granules with no halos and transforming granules with narrow halos) to the number of moles of crystalline insulin (in mature granules with wide halos), which was determined to be *Q*_*proins*_*/Q*_*ins*_ = 0.046 ± 0.005; (ii) half-life of insulin release from fusion of granules with the plasma membrane by pulse-chasing islets with ^35^S-labeled amino acids, which was determined to be τ_R_ = 96±12 hrs; and (iii) measured maturation time of isolated proinsulin-containing immature granules through action of prohormone convertases by pulse labeling with ^3^H-leucine, obtained from earlier work of Rhodes et al. (1987) and Furuta et al. (1998). Thus, as predicted, we find that *Q*_*proins*_*/Q*_*ins*_ ≈ τ_im_/τ_R_ based on our morphological measurements combined independent pulse-chase experiments.

Although we performed experiments on fixed specimens prepared for electron microscopy, without the ability to track subcellular dynamically in real time, our approach has some important advantages. First, the SBEM experiments were performed on beta cells in intact, cultured, wild-type pancreatic islets of Langerhans. Second, the SBEM experiments did not require the use of genetically-encoded, fluorescent labels to track secretory granules by optical microscopy, which can potentially introduce changes in the behavior of cultured cells (Ferri et al., 2019); the only labels used in the present experiments were isotopic derivatives of amino acids added to the islet incubation medium. Third, the SBEM experiments did not rely on immortalized insulinoma cell lines, which are widely used because of their ease of handling, but which are known to behave differently from beta cells in wild-type islets (Lightfoot et al., 2012).

### 3.5. Future work

Identification of secretory granule type in the mouse islet beta cells, as well as determination of the granule core volumes were performed manually by tracing the contours of about 8,000 mature granules and 800 immature or transforming granules, which is a laborious process, even though we analyzed only about 5% of the SBEM images that were acquired from 6 cells. Along with other laboratories, we have been developing automated procedures for segmenting cellular organelles using deep learning methods based on convolutional neural networks (Guay et al., 2018). The data generated in these experiments would provide a sensitive test for automated segmentation methods since subtle differences in morphology between immature granules with no halos and transforming granules with narrow halos must be reliably discriminated by the deep learning algorithms. Errors in identifying granule types would have a large effect on reliability of automated segmentation methods for estimation of the granule maturation time.

## 4. Summary

We have used SBEM to visualize and analyze volumes of complete beta cells in wild-type cultured mouse pancreatic islets of Langerhans with the aim of estimating the maturation times of secretory granules in beta cells containing proinsulin. It is assumed that the islets are in homeostasis, so that the mean maturation time can be determined from the total volume of proinsulin in the cores of immature secretory granules relative to the total volume of insulin in the cores of mature secretory granules, along with previous measurements of the release time for insulin by fusion of mature granules with the plasma membrane. It was found that the mean maturation time for the immature granules that bud off the trans-Golgi network and are characterized by the absence of halos is 135±14 minutes, and a further 130±17 minutes for the lifetime of the transforming vesicles, which are identified by their narrow halos. Further studies that combine electron microscopy techniques with biochemical assays could be used to learn more about the steps involved in granule maturation and in insulin release from beta cells. The techniques could also be used to investigate secretion processes in other types of cells if structural differences exist between immature and mature secretory granules.

## Acknowledgments

This work was supported by the intramural research programs of the National Institute of Biomedical Imaging and Biomedical Engineering, and the National Institute for Dental and Craniofacial Research at the National Institutes of Health in Bethesda, Maryland.

---

**Supplemental Video 1.** A typical wild type mouse beta cell in periphery of islet, containing region rich in Golgi and pre-immature granules shown at higher magnification.

